# Shallow metagenomics with Nanopore sequencing in canine fecal microbiota improved bacterial taxonomy and identified an *uncultured CrAssphage*

**DOI:** 10.1101/585067

**Authors:** Anna Cuscó, Anna Salas, Celina Torre, Olga Francino

## Abstract

Long-read metagenomics –using single-molecule sequencers– has the potential to assembly entire genomes, even from complex metagenomics datasets. Using long-read metagenomics with Nanopore sequencing in pooled samples, we aim to improve the individual taxonomic profiles obtained with V4 16S rRNA massive sequencing and to assemble the fecal metagenome of healthy dogs.

Fecal samples from healthy dogs were sequenced individually using V4 16S rRNA gene and in pools using a shallow metagenomics approach with Nanopore sequencing. Long-read metagenomics allowed us refining the V4 16S rRNA results up to the species level and determining the main bacterial species inhabiting on fecal microbiota of our cohort of healthy dogs. Among them, the most abundant were *Fusobacterium varium; Megamonas hypermegale; Bacteroides vulgatus; Blautia hansenii; Clostridium perfringens*; and *Clostridoides difficile*. Once performed the metagenome assembly, one contig was suggested to be circular and hit to an *uncultured crAssphage*.

To conclude, shallow long-read metagenomics with pooled samples using MinION allowed characterizing the dog fecal microbiota at lower taxonomic level, such as bacterial species. The assembly of the reads retrieved a contig that represents a circular draft genome of an uncultured *CrAssphage* from dog fecal samples that is one of the most abundant bacteriophages in the human gut.

## Introduction

Metagenomics allows observing microorganisms from the whole tree of life, going beyond bacteria. The most common approaches rely on shotgun metagenomics and massive sequencing technologies, which sample the DNA from a specific environment, fragment it and sequence the small fragments with the aim to solve the puzzle (1). Single molecule sequencing platforms such as those developed by PacBio and Oxford Nanopore Technologies allow sequencing with longer reads, despite presenting lower accuracy (2).

Long-read metagenomics allows easier assembly of entire genomes, even from complex metagenomics datasets. Nanopore sequencing combined with short-read polishing has allowed the assembly of complete or nearly complete circular genomes retrieved from several metagenomics samples. On one hand, this approach has been tested in bacterial (3,4) and viral mock communities (5). On the other hand, it has also been applied on real samples: i) to characterize human fecal metagenome (4,6,7); ii) to detect viruses on clinical samples (8); and iii) to characterize other environments, such as an enrichment reactor (9) or the cow rumen (10).

Until the date, two studies have characterized dog fecal microbiota using shotgun metagenomics, and both of them showed similarities to human gut microbiota (11,12). We aim to profile the canine fecal microbiota at lower taxonomical level -such as species-performing low deep metagenomics with Nanopore long-reads to complement the individual taxonomic profiles obtained with V4 16S rRNA sequencing. We performed the long-read metagenomics with pooled samples rather than individual ones to improve the cost/benefit ratio of our approach.

After assembling the fecal metagenome of healthy dogs, we retrieved several contigs, one of them suggested being circular and blasting to an *uncultured crAssphage*. *Uncultured crAssphage* and crAss–like bacteriophages are the most abundant bacteriophages in the human gut (13–16). This bacteriophage was discovered *in silico* in 2014 when screening multiple metagenomics sets (13). A crass-like bacteriophage was isolated for the first time in 2018 (17).

## Material and methods

### Sample collection and DNA extraction

Fecal samples were collected and frozen at −80°C until the DNA extraction. Bacterial DNA was extracted from 200 mg of fecal sample using the ZymoBIOMICS™ DNA Microprep Kit under manufacturer’s conditions. To assess for contamination from the laboratory or reagents, one blank sample was processed at the same time.

### V4 16S rRNA PCR, massive sequencing and bioinformatics analyses

V4 hypervariable region of 16S rRNA gene was amplified for each dog fecal sample using the Phusion Hot Start II High-Fidelity DNA polymerase (Thermo Scientific). The samples were amplified using the barcoded forward primer 515F and reverse primer R806, with sequencing adaptors at the 5′ end. One no template control (NTC) sample was included in each PCR reaction to assess if there was contamination. Each PCR reaction contained RNAse and DNAse free water, 5x Phusion Buffer HF (5 µl), dNTPs 2mM (2.5 µl), Primer Fw 10µM (1.25 µl), Primer Rv 10µM (1.75 µl), Phusion High Fidelity Taq Polymerase 2 U/µl (0.25 µl) and 5 ng of DNA. The thermal profile consisted of an initial denaturation of 30 sec at 98°C, followed by 30 cycles of 15 sec at °98 C, 15 sec at 50°C, 20 sec at 72°C, and a final extension of 7 min at 72°C.

Quality and quantity of PCR products were determined using Agilent Bioanalyser 2100 and Qubit™ fluorometer, respectively. Samples were sequenced on an Ion Torrent Personal Genome Machine (PGM) with the Ion 318 Chip Kit v2 (Life Technologies) under manufacturer’ conditions. Supplementary Table 1 contains information about the sequencing performance.

Demultiplexed fastq reads were analyzed using Quantitative Insight Into Microbial Ecology 2 (QIIME 2) software (18) (https://qiime2.org). DADA2 was used as quality filtering method to denoise, dereplicate single-end sequences and remove chimeras (19). Amplicon Sequence Varians (ASVs) were used to classify the sequences and assign taxonomy, using Greengenes 13.8 (20) at 99% of similarity to reduce redundancy of the database. Finally, chloroplasts were removed from the sequences. Taxonomic composition was plotted using ampvis2 package (21) from R software.

### Long read metagenomics, single molecule sequencing and bioinformatics analyses

Long read metagenomics was performed in two pools of DNA. Pool 1 compresses six fecal microbiota samples, while pool 2, three samples. Two samples from both pools come from the same dogs, but at two different time points. DNA samples were pooled considering the initial DNA quality and quantity to obtain an even representation of the samples included. Libraries were prepared with the Rapid Barcoding kit (SQK-RBK004) and sequenced using MinION, following the instructions of the Rapid Barcoding Sequencing protocol of Oxford Nanopore Technologies.

The samples were run using MinKNOWN software. Fast5 files generated by the software were basecalled and demultiplexed (sorted by barcode) using Albacore v2.3.1, obtaining fastq files. By default, sequences with a q-score lower than 7 are sent to fail folder. A second round of demultiplexing was done with Deepbinner (Wick et al., 2018) in which barcodes that agreed with Albacore were kept and the remaining ones were removed. Supplementary Table 1 contains information about the sequencing performance.

Reads shorter than 1,000 bp were removed for further analyses. To assign taxonomy we used *What’s in my Pot* (WIMP) (22), a workflow from EPI2ME in the Oxford Nanopore Technologies cloud (based on Centrifuge (23) software) that uses the NCBI RefSeq database. A final filtering step was applied and we kept only those hits with a classification score greater than 300 (23). Taxonomic composition was plotted using ampvis2 package (21) in R software.

Using the reads from both pools to obtain a sufficient number of sequences we performed the assembly using canu (24). We polished the contigs obtained using Nanopolish (25). We focused this first analysis on the predicted circular contig corresponding to an uncultured crAssphage. For this contig, Phaster (26) was used to predict the CDS. ORFs were blasted using pBLAST. Descriptions of the protein names were retrieved using UniProt (https://www.uniprot.org/uploadlists/). This contig was mapped using Minimap2 (27) against the crass-like genomes reported by Guerin and collaborators (14).

### Data availability

The datasets analyzed during the current study are available in the NCBI Sequence Read Archive, under the Bioproject accession number PRJNA527983.

## Results and discussion

### Fecal microbiota of healthy dogs

Fecal samples from healthy dogs were sequenced individually using V4 region amplicons of the 16S rRNA gene (massive sequencing) and in pools using a metagenomics approach with long-reads (single molecule sequencing by ONT). Results for both approaches are shown in Figure 1.

**Figure 1.**
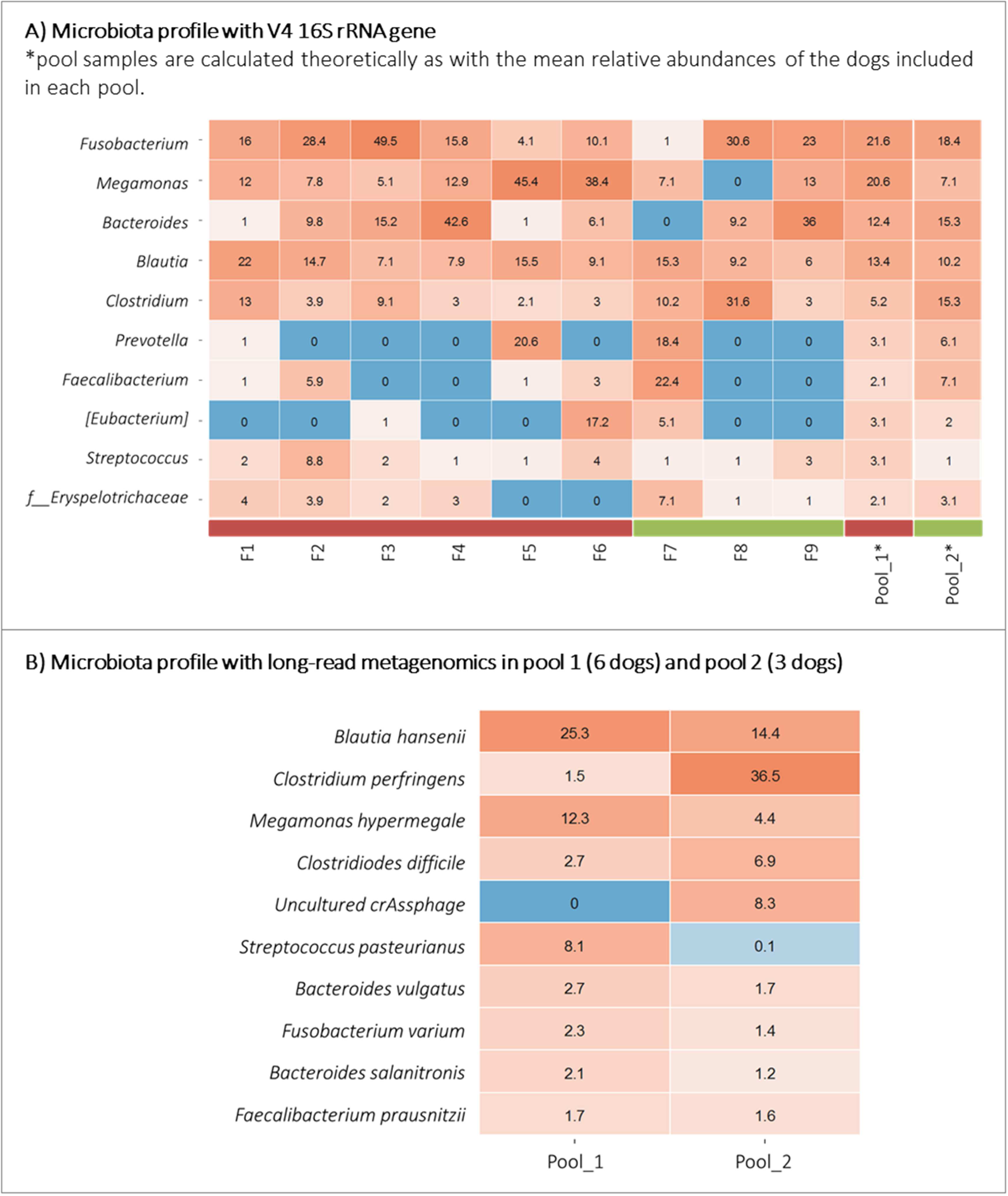
Heatmap representing fecal microbiota composition of healthy dogs. Each value represents the relative abundance of a specific taxa. A) Microbiota composition at the genus level as detected by V4 16S rRNA amplification and sequencing. The last two columns represent the theoretical mean value of each pool. B) Microbiota composition at the species level of pool 1 (6 dogs) and pool 2 (3 dogs) as detected by long-read metagenomics.

Results obtained for the 16S-V4 amplicon approach show that the fecal microbiota of healthy dogs from our cohort is composed by bacteria from the genera *Fusobacterium, Megamonas, Bacteroides, Blautia* and *Clostridium*, with relative abundance greater than 1% in each individual (except F7 and F8, for *Bacteroides* and *Megamonas* respectively). Other genera with lower abundance are *Prevotella, Faecalibacterium, Eubacterium, Streptococcus* and the family Eryspelotrichaceae.

We can refine microbial profiles retrieved by sequencing short fragments of the 16S rRNA approach down to species level either using long-amplicons of 16S rRNA gene or the 16S-ITS-23S amplicons (28) or using shallow long-read metagenomics. Here we applied the second approach in pools of samples. We pooled the DNA samples that had been previously used for the 16S rRNA gene amplification and short-read sequencing (V4) and we sequenced them with MinION device from ONT in a metagenomics approach. After quality filtering and basecalling, we assigned taxonomy with WIMP (22).

The results obtained in this long-read metagenomics approach revealed that the main bacterial species on the fecal microbiota of this cohort of dogs are: *Fusobacterium varium; Megamonas hypermegale; Bacteroides vulgatus; Blautia hansenii*; and *Clostridium perfringens*, together with *Clostridoides difficile*, as representatives of *Clostridium* genus. Other species are *Streptococcus pasteurianus, Bacteroides salanitronis and Faecalibacterium prausnitzii* (see Figure 1B).

For bacterial genomes that present similar genome sizes, WIMP could be considered a semi-quantitative approach. Even though, we observed some divergences on the relative abundances when comparing V4 16S rRNA gene approach and long-read metagenomics. Some of the divergences could also be due to the variable number of 16S rRNA gene copies depending on the bacterial species (29) or to the bias associated with the universal primers chosen for the amplification (30).

The metagenomics approach allowed us detecting also non-bacterial members of the fecal microbiota. The most abundant one was *uncultured crAssphage*, representing around 8% of total relative abundance on Pool 2. *CrAssphage* infect bacteria from the *Bacteroides* genus and other Bacteroidetes members (13). Moreover, the first study culturing a *crAssphage* (ΦCrAss001) from human fecal samples showed that it infected *Bacteroides intestinalis* (17).

Previous studies on dog fecal shotgun metagenomics suggested that the canine gut microbiota resembles that from humans (11); more recently, Coelho and collaborators showed that canine gut microbiota resembles more to human gut microbiota than the most commonly used animal models, such as pig or mouse (12). Our results suggest another similarity: *uncultured crAssphage* was the most abundant virus in the fecal microbiota of one cohort of healthy dogs, similar to what has been previously reported in humans (13–15).

### Assembly of an uncultured *crAssphage* from canine fecal metagenome

Long-read metagenomics allows easier assembly of entire genomes, even from complex metagenomics datasets, as it has been seen in previous studies (3–10). We used the long reads obtained by nanopore sequencing in the metagenomics approach to perform assembly and polishing. Using this strategy we were able to retrieve several contigs. One of the contigs with 107,597 bp and coverage of 6.8x was suggested to be circular, and it blasted to uncultured *crAssphage* (NC_024711.1) with a query coverage of 96% and identity of 96.72% (Figure 2 and Supplementary Table 2). We also aligned our contig with the crass-like genomes from Guerin and collaborators (14) and it mapped with the highest score to ERR844050_ms_1 and to ERR844032_ms_1. Both genomes belong to the candidate AlphacrAssvirinae subfamily that contain the prototypical *crAssphage* (Supplementary Table 2) and present an identity greater than 97% to the reference crAssphage genome (NC_024711.1).

**Figure 2.**
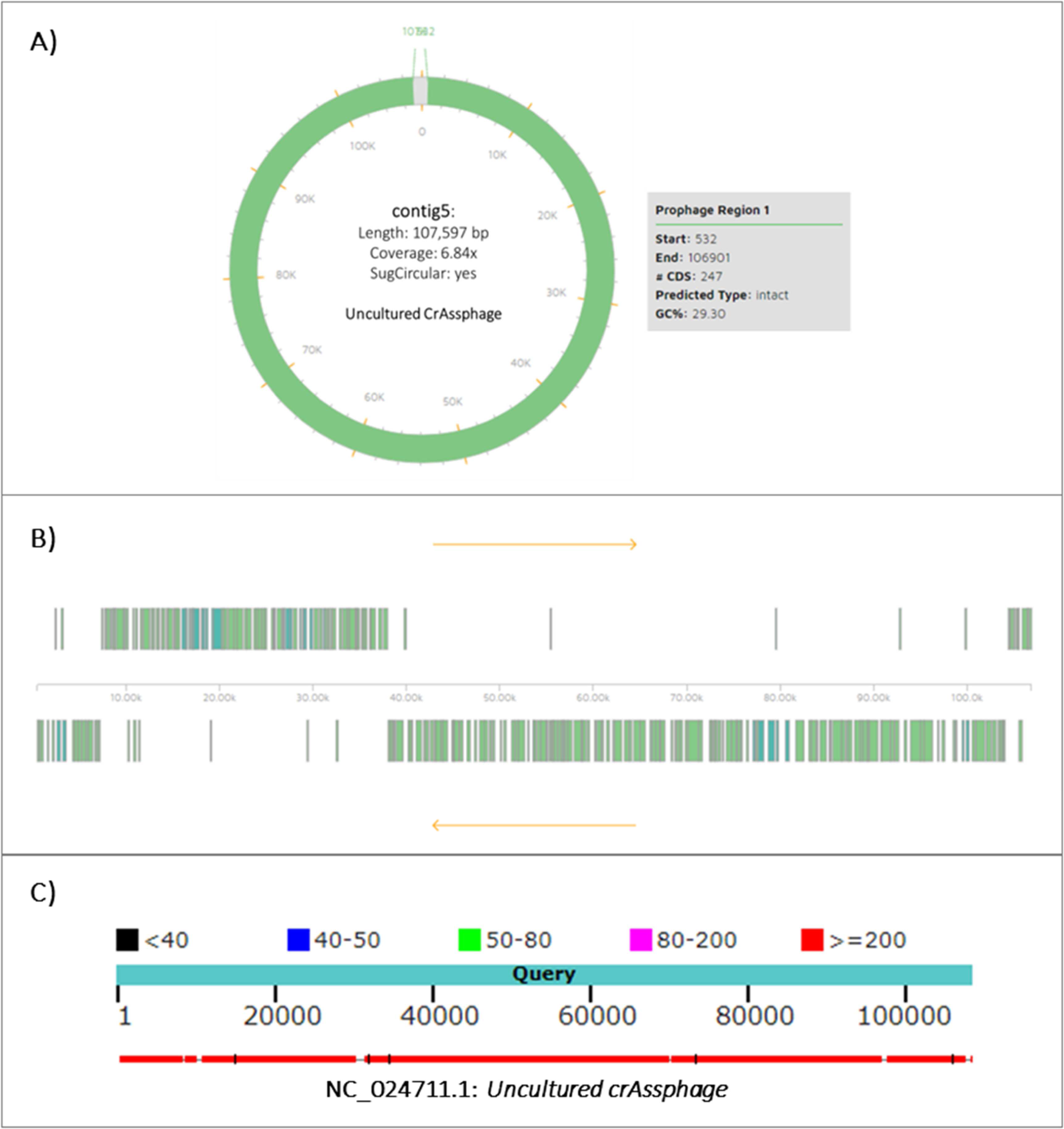
*Uncultured crAssphage* contig. A) Graphical representation of the uncultured *crAssphage* contig retrieved by Phaster, with some summary statistics associated from the assembly with canu. B) Genome representation of the *uncultured crAssphage* contig: green represents hypothetical proteins, and blue represents phage-like proteins as predicted by Phaster (see the detailed ORF predictions in Supplementary file 3). C) BLAST alignment with the Refseq database to *Uncultured crAssphage* (NC_024711.1), Query coverage of 96% and identity of 96.42% (see the complete alignment BLAST results in Supplementary File 2).

The NCBI reference genome (NC_024711.1) for *uncultured CrAssphage* contained 88 CDS, while preliminary results with Phaster (26) predicted a total of 247 on our contig (Supplementary Table 3). Long-read sequencing technologies allow assembling draft genomes from metagenomes, but they present insertion/deletion errors that affect protein prediction, often truncating them (31), as seen here. This draft assembly will be further improved when combining with short reads in a hybrid assembly approach.

Even having an unpolished genome, we were able to identify the two main genome regions with opposite gene orientation of crAss-like phages (Figure 2B)(14,15) and their associated proteins despite being truncated –splitted in several CDS (Supplementary Table 3). The small region contains proteins involved in replication, while the bigger region encodes for proteins related to transcription and virion assembly. The predicted proteins were found in the same order as previously reported for *CrAssphage* (Supplementary Table 3) (14,15). Further polishing using short reads would allow a better characterization of the *crAssphage* genome found in canine fecal metagenome.

*Uncultured crAssphage* was the most abundant virus previously reported in human fecal metagenomes (13–15) and it is globally spread throughout the industrialized societies (32,33). Since the initial screening of *CrAssphage* via PCR in several animal cohorts was negative, it has been regarded as a potential marker for human fecal pollution, to distinguish the origin of the pollution from other animal sources (34,35). Using this methods, dog fecal samples have already been screened for *CrAssphage*, with negative results (35,36). However, our results suggest that uncultured *crAssphage* can be an abundant member of dog fecal metagenome.

Our results would be in line with previous articles that have detected *crAssphage* via qPCR in fecal samples from cats, and wastewater derived from cattle, pigs and poultry (36,37). Since these studies were PCR-based, they could not discard the possibility of a cross-reaction of the PCR to some similar genomic regions of other related phages. CrAssphage has also been detected in one Gorilla that had a close contact with humans (32). Here, we obtained a circular contig identified as an *uncultured crAssphage* and it presented a high average nucleotide identity with *crAssphage* genomes isolated in human feces (Supplementary Table 2). Larger studies screening more dogs and other mammals that have close contact with humans should be performed to confirm our preliminary results and to assess the potential of *crAssphage* to be used as a biomarker of fecal pollution from either human or other species origin.

In conclusion, shallow long-read metagenomics with pooled samples using MinION from Oxford Nanopore Technologies allowed characterizing the dog fecal microbiota at lower taxonomic level, such as bacterial species. The assembly of the reads retrieved a contig that represents a circular draft genome of an uncultured *CrAssphage* from dog fecal samples. Hybrid assembly using shorter reads from massive sequencing technologies will allow a better characterization of the *CrAssphage* found in dogs and approaches including Hi-C could allow determining the host of this bacteriophage in dogs.

## Supporting information

Run summary

Blast and mapping results

Phaster output

